# Differential cortical and thalamic engagement of cholinergic interneurons in the nucleus accumbens core

**DOI:** 10.64898/2026.03.06.710127

**Authors:** Emily V. Jang, Adam G. Carter

## Abstract

Cholinergic interneurons (CINs) in the nucleus accumbens (NAc) play a key role in regulating motivated behaviors. Here we examine the connectivity and functional impact of cortical and thalamic inputs onto CINs in the NAc core. We first use cell-type specific retrograde anatomy to identify the prefrontal cortex and thalamus as putative afferents. We then combine *ex vivo* slice physiology and optogenetics to characterize the properties of synapses onto CINs. We demonstrate that thalamic inputs strongly facilitate, whereas cortical inputs exhibit marked depression. We also show that a combination of AMPA and NMDA receptors contribute to both cortical and thalamic responses. Lastly, we establish how these inputs and receptors evoke action potentials and influence spontaneous firing. Our findings show how CINs in the NAc core process long-range inputs, highlighting differences from equivalent circuits in other parts of striatum.

**SIGNFICANCE STATEMENT:** Cholinergic interneurons provide the primary source of acetylcholine in the striatum and are important for behavior and disease. The types of afferents that drive these interneurons have been examined in dorsal striatum but remain understudied in the nucleus accumbens. We found that inputs from prefrontal cortex and thalamus are the main drivers in the mouse nucleus accumbens core. We compare the sign, dynamics, and impact of these two excitatory inputs, showing how they engage multiple glutamate receptors to influence cholinergic interneuron firing.

## INTRODUCTION

The nucleus accumbens (NAc) is part of the ventral striatum that helps orchestrate motivated behaviors (Mogenson et al., 1980; Humphries and Prescott, 2010; Haber, 2011; Mannella et al., 2013; Floresco, 2015). The NAc consists of core and shell subregions (Záborszky et al., 1985; Zahm and Brog, 1992; Brog et al., 1993), which process a variety of inputs from cortical, thalamic, and limbic areas (Groenewegen et al., 1999; Smith et al., 2004; Ambroggi et al., 2008; Sesack and Grace, 2010; Britt et al., 2012). These afferents engage multiple cell types, including medium spiny neurons (MSNs) and local interneurons (MacAskill et al., 2012; Li et al., 2018; Scudder et al., 2018; Baimel et al., 2019). Cholinergic interneurons (CINs) are a small fraction of cells in the NAc (Wilson et al., 1990; Kawaguchi et al., 1995; Tepper and Bolam, 2004; Muñoz-Manchado et al., 2018), and responsible for releasing acetylcholine that shapes MSN firing (Witten et al., 2010) and dopamine release (Cachope et al., 2012; Yorgason et al., 2017). However, the ability of different long-range inputs to influence the firing of CINs in the NAc core is largely unexplored.

CINs have been primarily studied in the dorsal striatum, where they receive inputs predominantly from cortex and thalamus, which have distinct presynaptic and postsynaptic properties (Smeal et al., 2008; Ding et al., 2010; Doig et al., 2014; Guo et al., 2015; Johansson and Silberberg, 2020). Thalamic inputs derive primarily from the parafascicular nuclei, exhibit pronounced synaptic facilitation, and strongly drive CIN firing (Lapper and Bolam, 1992; Ding et al., 2010; Doig et al., 2014). Inputs from sensorimotor cortex have less influence on CIN firing and have been found to display either modest facilitation (Ding et al., 2010; Johansson and Silberberg, 2020) or depression (Doig et al., 2014; Mamaligas et al., 2019). Synaptic responses can be mediated by both AMPA and NMDA receptors (AMPAR and NMDAR) (Johansson and Silberberg, 2020), although NMDARs are more engaged by thalamic inputs (Kosillo et al., 2016; Mamaligas et al., 2019). The NAc also processes cortical and thalamic inputs, but whether the synaptic connectivity and physiology of CINs in the NAc core are equivalent to dorsal striatum remains unknown.

Connectivity within the NAc strongly depends on location, with nearby areas often receiving very different long-range afferents. The NAc medial shell receives inputs from the ventral hippocampus, basolateral amygdala, and paraventricular thalamus (Phillipson and Griffiths, 1985; Berendse and Groenewegen, 1990; Britt et al., 2012), and CINs in this subregion mostly process hippocampal and thalamic inputs (Baimel et al., 2022). In contrast, NAc core receives inputs from the prefrontal cortex and medial and midline thalamus, with few inputs from ventral hippocampus (Berendse and Groenewegen, 1990; Brog et al., 1993; Groenewegen et al., 1999; Li et al., 2018). These long-range inputs are known to make contact with MSNs in the NAc core (MacAskill et al., 2012), but their influences on CINs has not been examined. One possibility is that the NAc core resembles the dorsal striatum, with thalamus dominating the activation of CINs. Another possibility is that the prefrontal cortex serves as a dominant input, equivalent to hippocampal inputs to the NAc medial shell. Establishing the main drivers of CINs is important, as these interneurons ultimately exert a strong influence on the local network and help to shape behavior.

Here we use cell-type specific retrograde anatomy, *ex vivo* electrophysiology, and optogenetics to examine how CINs in mouse NAc core process long-range excitatory inputs. We first establish that CINs receive inputs predominantly from the prefrontal cortex and thalamus, rather than ventral hippocampus or basolateral amygdala. We show that while both cortical and thalamic inputs are excitatory, they display distinct short-term dynamics, with more facilitation at thalamic inputs. Nevertheless, we find that both inputs activate both AMPARs and NMDARs to drive pronounced firing of CINs from either quiescent or firing states. Together, our results highlight the unique synaptic properties of cortical and thalamic afferents to CINs in the NAc core. These inputs are distinct from equivalent connections in both the dorsal striatum and NAc medial shell, with important implications for how cholinergic interneurons can be engaged by other brain regions.

## MATERIALS AND METHODS

### Animals

All experiments were approved by and conducted in accordance with the guidelines of the Institutional Animal Care and Use Committee (IACUC) at New York University. ChAT-eGFP (Jackson Laboratories, stock # 007902) and ChAT-IRES-Cre (Jackson Laboratories, stock # 006410) mice were crossed with wild-type C57BL/6J mice (originally purchased from Jackson Laboratories). Male and female animals were used for all experiments. Mice were group-housed with same-sex littermates under a 12-hour light-dark cycle with food and water given *ad libitum*.

### Stereotaxic injections

Stereotaxic injections were performed on P42–P70 mice as previously described (MacAskill et al., 2012, 2014; Scudder et al., 2018; Baimel et al., 2019, 2022). Mice were anesthetized by inhalation of isoflurane-oxygen mixture and head-fixed in a stereotaxic frame (Kopf). A burr hole was made over the injection site, through which viruses were injected. Injection site coordinates were as follows relative to bregma (in mm): NAc core = ML -2.2, DV -3.7, AP +2.1 at 18° angle; mPFC = ML -0.35, DV -2.3 and -2.1, AP +2.3 at 0° angle; mTH = ML -1.1, DV -3.7 and -3.5, AP –0.2 at 15° angle) (Franklin & Paxinos 3^rd^ edition). Only the right hemisphere of the brain was used. Borosilicate pipettes were backfilled and pressure-injected at 20 nL/sec into the target site using Nanoject III (Drummond) with 30-45 second inter-injection intervals. Pipettes were left in place for 10 minutes before being slowly retracted. After all injections, animals were given ketoprofen (5 mg/kg) before being returned to their home cages.

For monosynaptic rabies tracing, helper viruses AAV1-EF1a-FLEX-TVA-Cherry (180 nL) (UNC Vector Core) and AAV9-CAG-FLEX-oG (270 nL) (Salk) were unilaterally injected into the dorsomedial side of the NAc core in ChAT-IRES-Cre mice aged P42–P56. After allowing 5 weeks for expression, pseudotyped rabies virus SADΔG-GFP(EnvA) (470 nL) (Salk) was injected into the same location. Mice were then housed for 7 days before being perfused for anatomy. For anterograde anatomy and physiology experiments examining inputs onto CINs, AAV1-hSyn-hChR2(H134R)-EYFP (360 nL) (Addgene, 26973-AAV1) or AAV1-CaMKII-hChR2(H134R)-mCherry (360 nL) (Addgene, 26975-AAV1) was injected into mPFC or mTH of ChAT-eGFP. Mice were then housed for 16-21 days before being prepared for anatomy or *ex vivo* recordings.

### Slice preparation

Following isoflurane-induced anesthesia, virally injected mice were perfused intracardially with an ice-cold cutting solution containing the following (in mM): 65 sucrose, 76 NaCl, 25 NaHCO_3_, 1.4 NaH_2_PO_4_, 25 glucose, 2.5 KCl, 7 MgCl_2_, 0.4 Na-ascorbate, and 2 Na-pyruvate (bubbled with 95% O_2_/5% CO_2_). 300 µm coronal sections were cut in this solution and transferred to a holding chamber with artificial cerebral spinal fluid (ACSF) containing the following (in mM): 120 NaCl, 25 NaHCO_3_, 1.4 NaH_2_PO_4_, 21 glucose, 2.5 KCl, 2 CaCl_2_, 1 MgCl_2_, 0.4 Na-ascorbate, and 2 Na-pyruvate (bubbled with 95% O_2_/5% CO_2_). Slices recovered in ACSF for an additional 30 minutes in a 35°C water bath before being stored for at least 30 minutes at 24°C prior to recording. All experiments were conducted in ACSF held at 30-32°C.

### Electrophysiology

Whole-cell patch clamp recordings were made in ChAT-eGFP mice from fluorescently identified CINs in the rostral NAc core (between 1.70 and 1.10 mm anteriorly from bregma). For sequentially paired recordings, putative MSNs were identified by a lack of fluorescence in ChAT-eGFP mice. The NAc core was identified as the area around the anterior commissure, between the NAc medial shell and dorsal striatum. Cells were visualized using differential interference contrast (DIC) optics on an upright microscope (Olympus BX51) equipped with a CMOS camera (Hamamatsu).

For voltage-clamp experiments, borosilicate pipettes (3-5 MΩ) were filled with a Cs-based internal (in mM): 130 Cs-gluconate, 10 HEPES, 10 Na-phosphocreatine, 4 Mg_2_-ATP, 0.4 NaGTP, 10 TEA, 2 QX-314, and 10 EGTA, pH 7.3 with CsOH. For current-clamp recordings, pipettes were filled with a K-based internal (in mM): 135 K-gluconate, 7 KCl, 10 HEPES, 10 Na-phosphocreatine, 4 Mg_2_-ATP, 0.3 NaGTP, and 0.5 EGTA, pH 7.3 with KOH. For biocytin-filled cells, biocytin-HCl (5 mg/mL) (Tocris) was added to the K-gluconate internal. In some recordings, Gabazine (GZ) (10 µM), CPP (10 µM), and NBQX (10 µM) were used to block GABA_A_R, NMDAR, or AMPAR-mediated responses, respectively. To record monosynaptic inputs, TTX (1 µM), 4-AP (100 µM), CPP (10 µM) and extra Ca^2+^ (4 mM) were added to the bath. No synaptic blockers were included for recordings of intrinsic properties. Reversal potentials for AMPAR- and GABA_A_R-mediated currents were experimentally determined for CINs and MSNs.

In current-clamp recordings, current was injected to either maintain a resting membrane potential close to -50 mV or sufficiently depolarize the CINs to fire 1-2 Hz at baseline. Baseline firing rate was defined as the average frequency in the 2 seconds before stimulus delivery, and cells that could not be maintained at a 1-2 Hz baseline firing rate were excluded from analysis.

Electrophysiological data were collected with a MultiClamp 700B amplifier (Axon Instruments) and National Instruments boards using custom software in MATLAB (MathWorks). Signals were sampled at 10 kHz and filtered at 2 kHz for voltage-clamp and 5 kHz for current-clamp recordings. Series resistance was monitored to be less than 25 MΩ and not compensated.

### Optogenetics

Neurotransmitter release was triggered by activating ChR2 present in presynaptic terminals of mPFC or mTH in the NAc core, as previously described (MacAskill et al., 2012, 2014). Optical stimulation was obtained with 2 ms pulses of 470 nm light from a blue LED (Thorlabs). Experiments for isolating monosynaptic inputs with TTX and 4-AP used optical stimulation through a 60X immersion objective (Olympus) with a power range of 2-10 mW, as measured at the back of the focal plane of the objective, to elicit graded responses. For other recordings of CINs, LED stimulation was delivered through a 10X 0.3 NA immersion objective (Olympus) at one stimulus intensity (10 mW). In voltage-clamp recordings, sequential sweeps of 1, 2, 3, 4, and 5 pulses at 20 Hz were delivered. In current clamp recordings, 5 pulses of 10 mW optogenetic stimulation at 20 Hz were used. Inter-trial interval for single pulse stimulation was 10 seconds, and up to 25 seconds for trains.

### Histology and image analysis

Mice were anesthetized and perfused intracardially with 0.01 M PBS followed by 4% PFA. Brains were stored in 4% PFA for 12-24 hours at 4°C before being washed three times in 0.01 M PBS. 70 µm-thick slices were cut on a VT-1000S vibratome (Leica) and directly mounted onto gel-coated glass slides and cover-slipped using VectaShield with DAPI (Vector Labs). For biocytin-filled reconstructions of individual cells, slices were transferred from the recording chamber into 4% PFA for 12-24 hours at 4°C. Slices were incubated with streptavidin-647 (1:1000 in 0.01 M PBS) and 5% triton-x100 before being mounted for imaging. Fluorescent images were taken on an Olympus VS120 microscope using a 10X 0.25NA objective, or a Leica TCS SP8 confocal microscope using a 20X 0.75NA or 40X 1.3NA objective. All images were processed using Fiji.

### Abbreviation of mouse brain structures

All mouse brain structures are referenced to the Allen Reference Atlas (atlas.brain-map.org). Cortex abbreviations: ACC, anterior cingulate; PL, prelimbic; IL, infralimbic; ORBm, medial orbital; ORBl, lateral orbital; ORBvl, ventrolateral orbital; AI, agranular insular; RSP, retrosplenial; SSp, primary somatosensory; MOp, primary motor; MOs, secondary motor. Thalamus abbreviations: AM, anteromedial; AD, anterodorsal; IAD, interanterodorsal; IMD, intermediodorsal; MD, mediodorsal; PVT, paraventricular; PT, parataenial; RE, reuniens; CM, central medial; CL, central lateral; PF, parafascicular; VM, ventromedial. Other abbreviations: VTA, ventral tegmental area; BNST, bed nucleus stria terminalis; GP, globus pallidum; VP, ventral pallidum.

### Quantification and statistical analysis

All electrophysiology and anatomical data were replicated in at least 3 animals. Experimenters were not blind to experimental groups. No pre-test analyses were used to estimate sample sizes.

For anatomical experiments examining monosynaptic inputs, brain slices were aligned to the Allen Common Coordinate Framework (Wang et al., 2020). Cell bodies were manually counted for slices from rostral-caudal coordinates +2.6 to -4.8 relative to bregma. Animals with more than 30% labeled starter cells outside the NAc core were excluded from the final dataset.

Electrophysiology data were analyzed with Igor Pro (WaveMetrics). EPSCs and IPSCs amplitudes were averaged in a 1 ms window around the peak. For stimulation trains, amplitudes of pulses 2 through 5 were measured by subtracting the decay of the prior pulse from the peak. The paired pulse ratio was calculated as PPR = EPSC_n_ / EPSC_1_, where n = pulse number. Synaptic charge was calculated as total area under the EPSC (pA * ms). For stimulation trains, the charge contribution of pulse (n) was calculated by subtracting the charge of pulse (n-1) from the cumulative charge. In the figures, average traces of synaptic currents show mean ± SEM. For current-clamp recordings, spikes were grouped in either 10 or 50 ms bins to calculate either the probability of firing or the firing rate. Statistical significance of differences was evaluated using the Wilcoxon Signed-Rank test for paired data, the Friedman test with Dunn’s multiple comparisons test for data with multiple time points, or with repeated measure 2-way ANOVA followed by Tukey or Sidak’s multiple comparison with p values < 0.05 considered significant. Statistical tests and graph generation were performed using Prism 9 (GraphPad).

## RESULTS

### Characterization of cholinergic interneurons

We used ChAT-eGFP mice to identify CINs by their green fluorescence, observing a sparse distribution in the NAc core (**Fig. 1A**). To characterize their intrinsic membrane properties and action potential firing, we performed whole-cell recordings from labeled cells in *ex vivo* slices (**Fig. 1B**). We determined that CINs in the NAc core exhibited little spontaneous firing (mean frequency = 0.28 ± 0.17 Hz; n = 12 cells / 6 mice) but could be reliably driven to spike with depolarization (**Fig. 1C**). CINs also displayed depolarized resting potentials (V_rest_ = -49.5 ± 1.4 mV), high input resistances (R_in_ = 257 ± 22 MΩ) and prominent voltage sag in response to hyperpolarization (sag = -9.6 ± 0.7 mV) indicative of *I*_h_ conductances (Kawaguchi, 1992) (**Fig. 1D**). These intrinsic membrane properties are similar to previous reports in the NAc core (Gallo et al., 2022), NAc medial shell (Witten et al., 2010; Baimel et al., 2022), and dorsal striatum (Straub et al., 2014), and indicate that CINs are excitable and rest close to threshold for firing.

**Figure 1:**
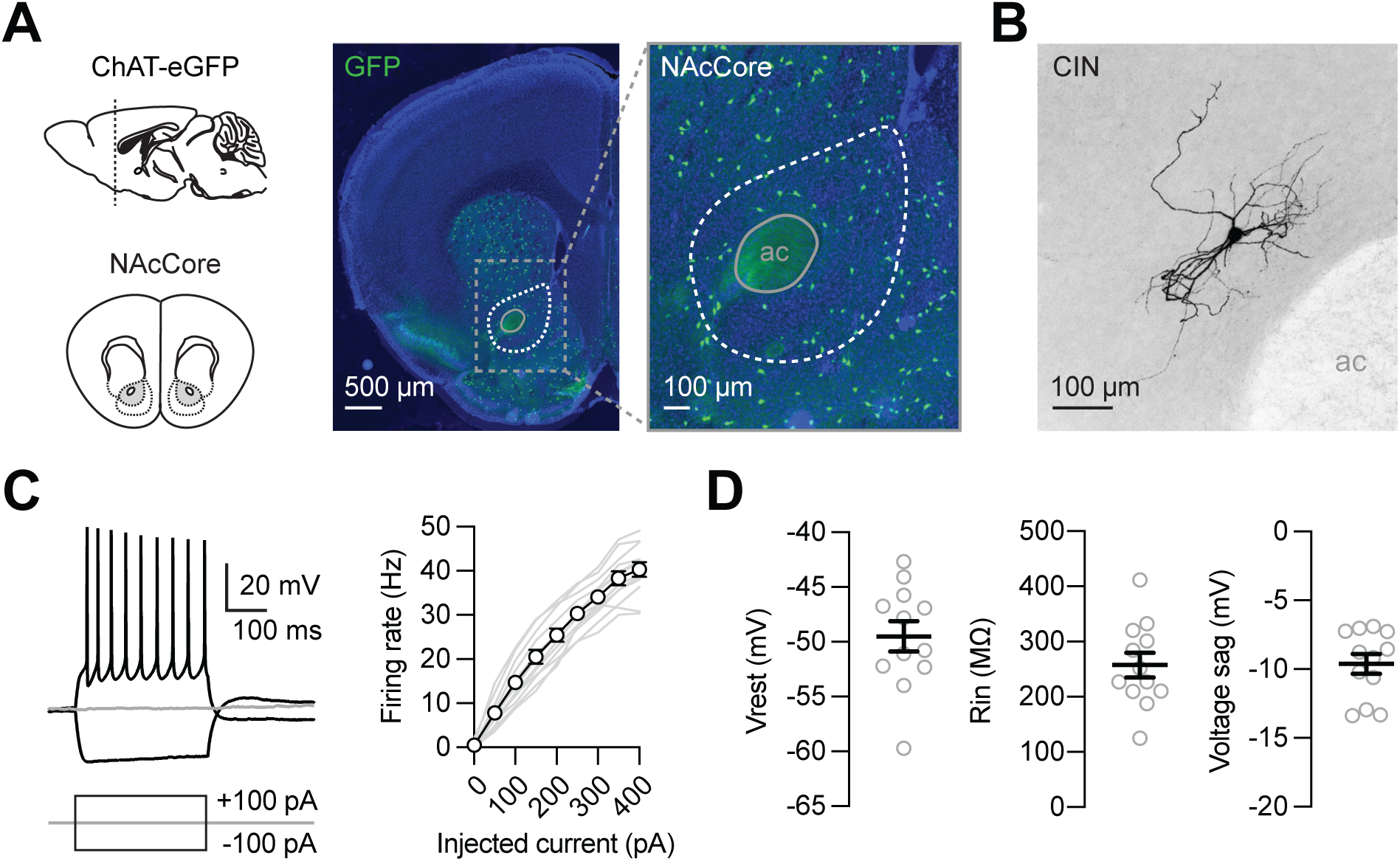
Characterization of cholinergic interneurons. **(A)** *Left,* ChAT-eGFP mice were used to identify cholinergic interneurons (CINs) in the NAc core. *Middle*, Coronal section showing the distribution of green CINs across the striatum. *Right,* Expanded view of NAc core (dotted white line) surrounding the anterior commissure (ac). **(B)** Representative confocal image of a biocytin-filled CIN in the NAc core. **(C)** *Left,* Representative current-clamp recording showing voltage response to current steps of -100, 0, and +100 pA. *Right,* Summary of firing rate vs. injected current curve. Gray lines are individual cells and black line is average ± SEM (n = 12 cells / 6 mice). **(D)** Summary of intrinsic membrane properties, including resting membrane potential (Vrest), input resistance (Rin), and voltage sag due to h-current (I_h_).

### Identification of long-range afferents

To survey the long-range inputs to CINs, we next used Cre-dependent monosynaptic rabies virus tracing (Wall et al., 2010). In ChAT-Cre mice, we first injected AAV1-FLEX-TVA-mCherry and AAV9-FLEX-oG helper viruses into the NAc core to target starter cells (n = 7 mice). Five weeks later, we then injected pseudotyped rabies virus EnvA-RV-GFP, which is taken up by TVA-expressing starter cells, restricting infection to CINs. The modified rabies virus spreads retrogradely from cells containing glycoprotein (oG), thus labeling monosynaptic inputs to CINs. After allowing one additional week for expression and transport, coronal sections across the entire brain were imaged and aligned to the Allen Institute’s Mouse Common Coordinate Framework (CCF) for a non-biased approach to manual cell-counting (Wang et al., 2020). Starter cells were identified by the co-expression of mCherry and GFP and found to be restricted to the injected area (**Fig. 2A**). Input cells expressing only GFP were found locally and across the brain (**Fig. 2B & Fig. S1**), with the majority mapped to cortical, thalamic, and pallidal regions (cortex = 29 ± 1%; thalamus = 28 ± 1%; pallidum = 24 ± 2% of long-range inputs) (**Fig. 2C & Fig. S1**). Cortical input cells originated from a variety of areas and particularly from frontal cortex (62 ± 4% of cortical inputs), including the medial prefrontal cortex (mPFC) (**Fig. 2D & Fig. S1**). Thalamic input cells primarily arose from medial and midline areas (medial = 37 ± 2%; midline = 27 ± 3% of thalamic inputs), which we refer together as mTH (**Fig. 2E & Fig. S1**). These anatomical data suggest that CINs in the NAc core likely receive long-range inputs from both prefrontal cortex and thalamus.

**Figure 2:**
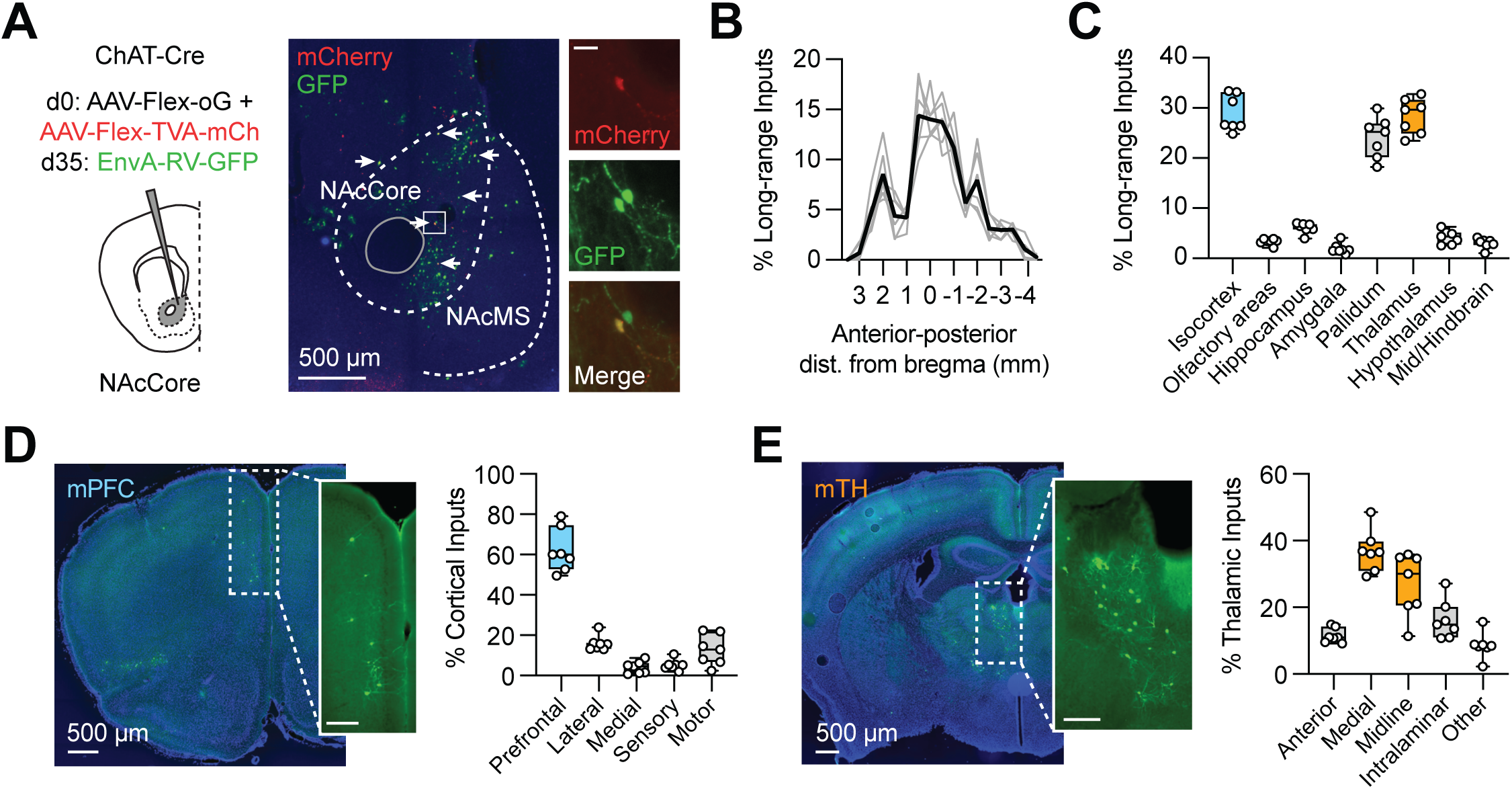
Identification of long-range afferents. **(A)** *Left,* Schematic showing injection of AAV-Flex-oG and AAV-Flex-TVA-mCherry into the NAc core of ChAT-Cre mice, followed 5 weeks later by EnvA-RV-GFP. *Middle,* Representative confocal image of injection site, indicating starter cells labeled with both mCherry and GFP (white arrows), surrounded by monosynaptic input cells labeled with only GFP. *Right,* Expanded views of starter and input cells in the NAc core (inset scale bar = 25 µm). **(B)** Distribution of all monosynaptically connected input cells located across the anterior-posterior axis relative to bregma (n = 7 mice). **(C)** Summary of long-range input cell locations, with cortex highlighted in blue and thalamus in orange. **(D)** *Left,* Example image of input cells in the medial prefrontal cortex (mPFC) (inset scale bar = 250 µm). *Right,* Distribution of cortical input cells by subregion, including prefrontal (62 ± 4%), lateral (16 ± 1%), medial (4 ± 1%), sensory (5 ± 1%), and motor (13 ± 3%). **(E)** *Left,* Example image of input cells in the medial and midline thalamus (mTH) (inset scale bar = 250 µm). *Right,* Distribution of thalamic input cells by subregion, including anterior (12 ± 1%), medial (37 ± 2%), midline (27 ± 3%), and intralaminar (16 ± 2%).

### Cortical and thalamic synaptic connections

Having identified possible long-range inputs to the NAc core, we next used slice electrophysiology to examine the functional properties of synapses onto CINs. We injected AAV1-hSyn-ChR2-GFP into the mPFC or mTH of ChAT-eGFP mice, waiting 2-3 weeks for expression and transport (**Fig. 3A & 3B)**. We observed that mPFC axons were present in the dorsal NAc core and dorsomedial striatum but largely absent in the nearby NAc medial shell (**Fig. 3A**). In contrast, mTH axons were found along the medial NAc core, parts of the NAc shell, and the dorsomedial striatum (**Fig. 3B**). These distributions suggest that although these long-range inputs exhibit distinct axonal profiles, they are well situated to make connections onto CINs in the NAc core.

**Figure 3:**
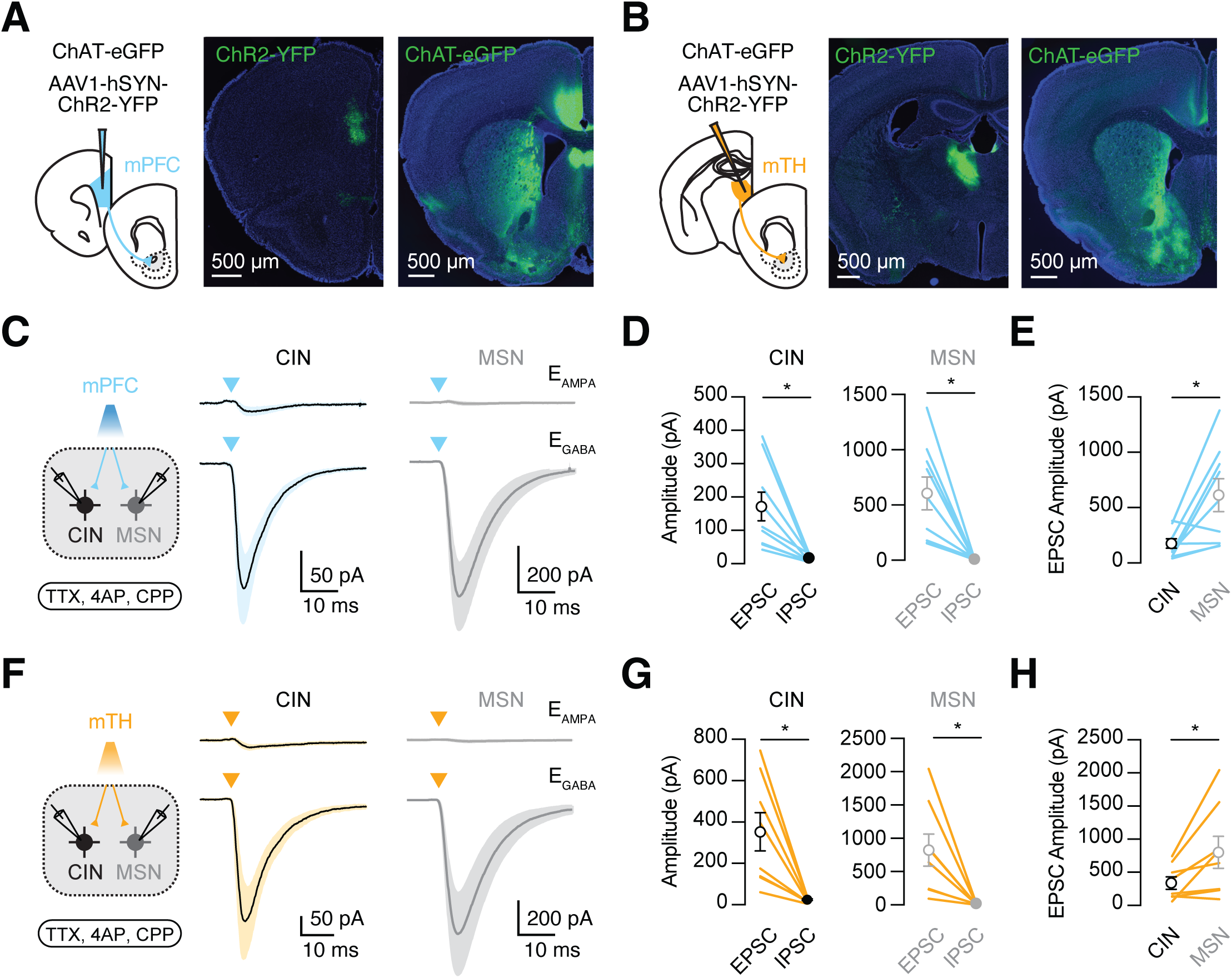
Cortical and thalamic synaptic connections. **(A)** *Left,* Schematic of AAV1-hSyn-mChR2-YFP injections into medial prefrontal cortex (mPFC) of ChAT-eGFP mice. *Middle,* Image of injection site. *Right,* Image of mPFC axons in striatum, including the NAc core. **(B)** Similar to (A) for injections into medial and midline thalamus (mTH). **(C)** mPFC-evoked EPSCs and IPSCs at nearby pairs of CINs (black) and MSNs (gray), recorded at reversal potentials for AMPAR (top) and GABA-A-R (bottom), respectively, in the presence of TTX, 4AP, and CPP (n = 9 cells / 5 mice). **(D)** Summary of mPFC-evoked EPSC and IPSC amplitudes, with lines showing individual CINs (left) and MSNs (right). **(E)** Summary of mPFC-evoked EPSC amplitudes at pairs of CINs and MSNs. **(F - H)** Similar to (C - E) for mTH inputs (n = 8 cells / 3 mice). * = P < 0.05

We next performed whole-cell voltage-clamp recordings from identified CINs, as well as nearby MSNs in the same slices. Comparing responses at neighboring cells allowed us to control for virus expression and light exposure (MacAskill et al., 2012). To isolate monosynaptic connections, we included TTX (1 µM), 4-AP (100 µM) and high Ca^2+^ (4 mM) in the bath, which blocks action potential propagation but restores presynaptic release (Petreanu et al., 2009). We also included CPP (10 µM) to block NMDARs, whose presence otherwise precludes isolating inhibitory postsynaptic currents (IPSCs) at positive holding potentials. We found that single mPFC inputs evoked prominent excitatory postsynaptic currents (EPSCs), but minimal IPSCs at CINs (EPSC = 168 ± 43 pA; IPSC = 14 ± 3 pA; EPSC vs. IPSC amplitude: Wilcoxon test: W = -45, p = 0.0039; n = 9 cells / 5 mice) (**Fig. 3C-E**). Similarly, we determined that single mTH inputs produced EPSCs but not IPSCs at CINs (EPSC = 351 ± 93 pA; IPSC = 21 ± 4 pA; EPSC vs. IPSC amplitude: Wilcoxon test: W = -36, p = 0.0078; n = 8 cells / 3 mice) (**Fig. 3F-H**). Both mPFC and mTH inputs elicited larger EPSCs at MSNs than CINs (mPFC evoked EPSCs, CIN vs. MSN: Wilcoxon test: W = 39, p = 0.0195; n = 9 pairs / 5 mice; mTH evoked EPSCs, CIN vs. MSN: Wilcoxon test: W = 34, p = 0.0156; n = 8 pairs / 3 mice) (**Fig. 3E & H**). However, neither mPFC nor mTH inputs elicited substantial IPSCs at MSNs, across light intensities (**Fig. S2**), indicating they are excitatory at both cell types (**Fig. 3C & F**). Together, these data indicate that cortical and thalamic inputs make robust excitatory connections onto CINs, where they may influence the firing properties of these cells.

### Short-term dynamics and receptor contributions

The strength of synaptic connections often varies with repetitive activity, in part due to changes in presynaptic release (Zucker and Regehr, 2002). At CINs in the dorsal striatum, short-term dynamics markedly differs between afferents, with more facilitation at thalamic inputs (Ding et al., 2010). To study equivalent dynamics in the NAc core, we next injected AAV1-CaMKII-ChR2-mCherry into mPFC or mTH of ChAT-eGFP mice (**Fig. S3**). In *ex vivo* slices, we delivered brief (2 ms) stimulus trains (1-5 pulses at 20 Hz) in the presence of gabazine (GZ) (10 µM) to block GABA-A receptors. To assess synaptic dynamics, we delivered incrementally increasing numbers of light stimuli, allowing us to subtract previous traces during trains. In voltage-clamp recordings, we found that trains of mPFC inputs (n = 8 cells / 6 mice) and mTH inputs (n = 8 cells / 5 mice) evoked robust responses at -50 mV and +40 mV (**Fig. 4A**). mPFC-evoked EPSCs were initially larger than mTH-evoked EPSCs at both -50 mV (mPFC EPSC_1_ = 158 ± 25 pA; mTH EPSC_1_ = 35 ± 9 pA; Mann Whitney test: U = 1; p = 0.0003) (**Fig. 4B**) and +40 mV (mPFC EPSC_1_ = 66 ± 15 pA; mTH EPSC_1_ = 21 ± 5 pA; Mann Whitney test: U = 10; p = 0.02) (**Fig. 4B**). However, with repetitive stimulation, the differences between these responses became non-significant at both -50 mV (mPFC EPSC_5_ = 158 ± 22 pA; mTH EPSC_5_ = 123 ± 12 pA; Mann Whitney test: U = 20; p = 0.23) (**Fig. 4B**) and +40 mV (mPFC EPSC_5_ = 122 ± 26 pA; mTH EPSC_5_ = 113 ± 14 pA; Mann Whitney test: U = 29; p = 0.80) (**Fig. 4B**). These data indicate that both mPFC and mTH axons evoke pronounced excitatory responses at CINs, but display distinct short-term dynamics, with more depression observed at cortical inputs and more facilitation at thalamic inputs.

**Figure 4:**
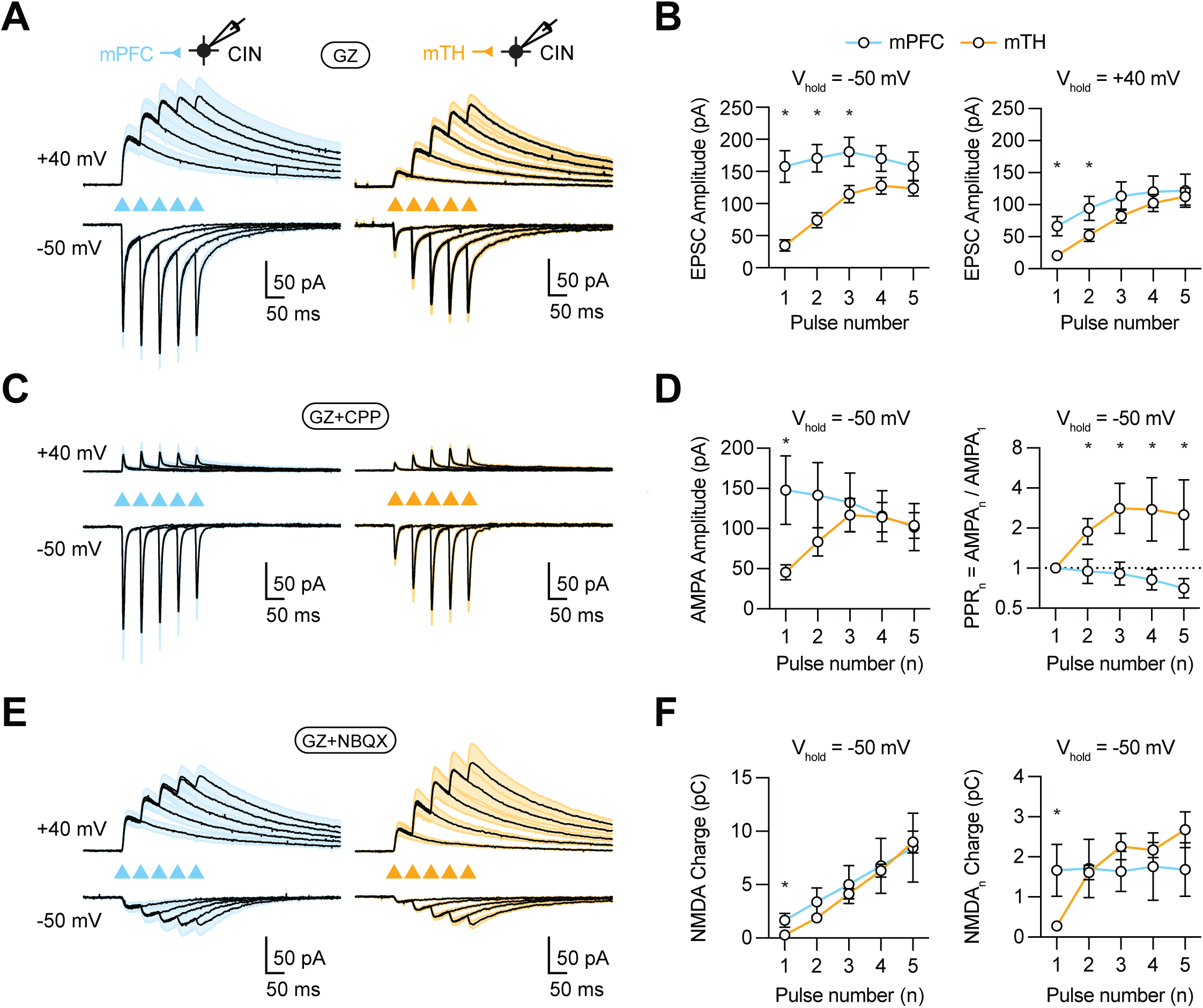
Short-term dynamics and receptor contributions. **(A)** Average mPFC (left) and mTH (right) evoked EPSCs at CINs in response to 1, 2, 3, 4 or 5 pulses of 20 Hz stimulation (traces overlaid) and recorded at +40 mV and -50 mV in the presence of gabazine (GZ) (mPFC: n = 8 cells / 6 mice; mTH: n = 8 cells / 5 mice). **(B)** *Left,* Summary of amplitudes of mPFC (blue) and mTH (orange) evoked EPSCs recorded at -50 mV as a function of pulse number. *Right,* Similar for ESPCs recorded at +40 mV. **(C)** Similar to (A) for AMPAR-mediated EPSCs evoked in the presence of GZ and CPP to block NMDARs (mPFC: n = 10 cells / 7 mice; mTH: n = 7 cells / 4 mice). **(D)** *Left,* Summary of amplitudes of mPFC (blue) and mTH (orange) evoked AMPAR-mediated EPSCs in CINs recorded at -50 mV as a function of pulse number (n). *Right,* Summary of paired-pulse ratio (PPR) calculated from these responses. **(E)** Similar to (A) for NMDAR-mediated EPSCs evoked in the presence of GZ and NBQX to block AMPARs (mPFC: n = 8 cells / 4 mice; mTH: n = 8 cells / 4 mice). **(F)** *Left,* Summary of cumulative charges of mPFC (blue) and mTH (orange) evoked NMDAR-mediated EPSCs in CINs recorded at -50 mV. *Right,* Charge as a function of pulse number (n), showing accumulation with each subsequent pulse. * = P < 0.05

To further explore these responses, we next performed two additional sets of recordings in the presence of either AMPAR or NMDAR antagonists. Using CCP (10 µM) to block NMDARs, we observed robust AMPAR mediated EPSCs at -50 mV evoked by both mPFC inputs (EPSC_1_ = 148 ± 43 pA; n = 10 cells / 7 mice) and mTH inputs (EPSC_1_ = 46 ± 9 pA; n = 7 cells / 4 mice) (**Fig. 4C**). As expected, responses were smaller at +40 mV, due to the smaller driving force for AMPAR conductances. Short-term dynamics again differed, with mPFC inputs showing depression (PPR_5_ = 0.7 ± 0.1) and mTH inputs facilitation (PPR_5_ = 3.0 ± 0.8; Mann Whitney test: U = 1, p = 0.0002) (**Fig. 4D**). Using NBQX (10 µM) to block AMPARs, we found robust NMDAR EPSCs in CINs at -50 mV evoked by mPFC inputs (n = 8 cells / 4 mice) and mTH inputs (n = 8 cells / 4 mice) (**Fig. 4E**). These responses were larger at +40 mV, as expected for relief of Mg2+ block at depolarized potentials. We quantified these responses as synaptic charge, which accumulated over trains to a similar degree for mPFC and mTH inputs, both at +40 mV (**Fig. S3**) and near the resting membrane potential of these cells (mPFC NMDAR charge = 8.5 ± 3.2 pC; mTH NMDAR charge = 9.0 ± 1.0 pC; Mann-Whitney test: U = 20; p = 0.23) (**Fig. 4F**). With CINs held at -50 mV, NMDAR EPSCs also displayed distinct dynamics for the two types of inputs (mPFC PPR_5_ = 1.1 ± 0.4, mTH PPR_5_ = 10.1 ± 1.5) (**Fig. 4F**). Together, these results indicate that both mPFC and mTH inputs engage AMPA and NMDA receptors but display distinct short-term dynamics.

### Functional impact on action potential firing

The properties of mPFC and mTH inputs suggest they may also be able to strongly impact action potential firing of CINs. To examine this influence, we next made current-clamp recordings from CINs at resting potentials in the presence of GZ (10 µM). In the dorsal striatum, CINs are predominantly driven by thalamic inputs, with less impact by cortical inputs (Kosillo et al., 2016; Mamaligas et al., 2019; Johansson and Silberberg, 2020). In contrast, we found that both mPFC and mTH inputs evoked robust firing of CINs in the NAc core (**Fig. 5A**). Consistent with our voltage-clamp recordings, mPFC but not mTH inputs evoked action potentials early in stimulus trains (probability of at least one spike in 10 ms after pulse 1: mPFC = 0.15 ± 0.08, n = 15 cells / 8 mice; mTH = 0.01 ± 0.00, n = 16 cells / 8 mice; Mann Whitney test: U = 76.5; p = 0.026) (**Fig. 5B**). However, in both cases, evoked action potentials built up across stimulus trains (probability of at least one spike in 10 ms after pulse 5: mPFC = 0.20 ± 0.06, n = 15 cells / 8 mice; mTH = 0.21 ± 0.04, n = 16 cells / 8 mice; Mann Whitney test: U = 105.5; p = 0.58) (**Fig. 5B**).

**Figure 5:**
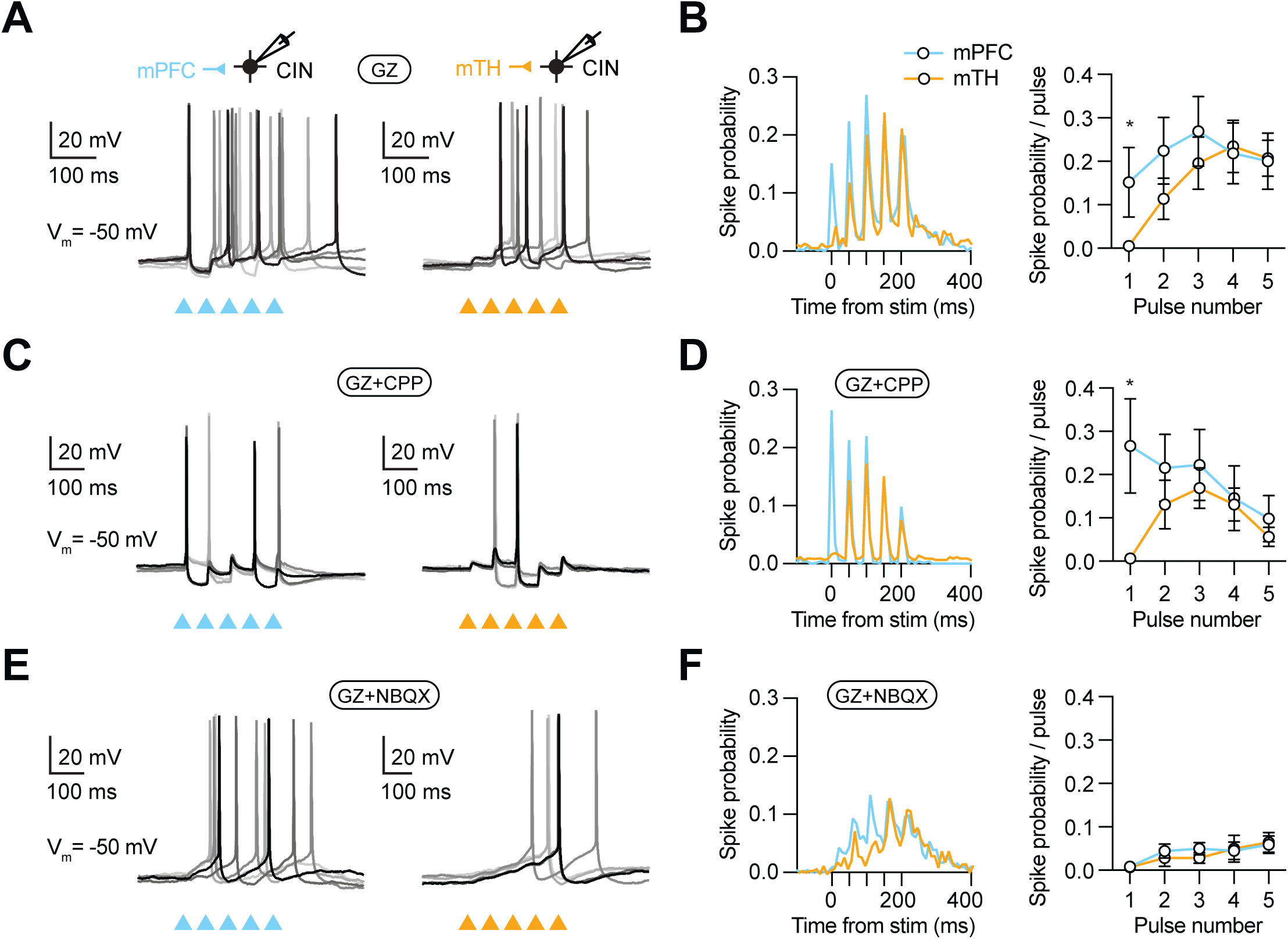
Functional impact on action potential firing. **(A)** Examples of action potentials evoked by mPFC inputs (left) or mTH inputs (right) in current-clamp recordings from quiescent CINs maintained close to -50 mV. **(B)** *Left,* Summary of spike probability versus time of stimulation for mPFC (blue) and mTH (orange) evoked firing (mPFC: n = 15 cells / 8 mice; mTH: n = 16 cells / 8 mice). Note that error bars are removed for clarity. *Right,* Summary of the probability of observing at least one action potential within 10 ms after each stimulation pulse. **(C - D)** Similar to (A - B) in the presence of GZ and CPP to isolate AMPAR evoked spikes (mPFC: n = 14 cells / 6 mice; mTH: 16 cells / 5 mice). **(E - F)** Similar to (A - B) in the presence of GZ and NBQX to isolate NMDAR evoked spikes (mPFC: n = 12 cells / 4 mice; mTH: n = 14 cells / 7 mice).

These findings indicate that despite different short-term dynamics, both cortical and thalamic inputs can robustly engage CINs. One possibility is that their influences on firing may reflect the effects of different types of glutamate receptors. To assess this influence, we next performed two sets of current-clamp recordings in the presence of selective receptor antagonists. In the presence of CPP (10 µM) to block NMDARs, both mPFC and mTH inputs evoked time-locked firing (**Fig. 5C & Fig. S4**). In this case, cortex-evoked responses depressed whereas thalamus-evoked responses facilitated (maximum spike probability for mPFC = pulse 1: 0.27 ± 0.09, n = 14 cells / 6 mice; vs. maximum spike probability for mTH = pulse 3: 0.17 ± 0.04, n = 16 cells / 6 mice) (**Fig. 5D**). In contrast, in the presence of NBQX (10 µM) to block AMPARs, both mPFC and mTH inputs evoked action potentials less time-locked to the stimulus (**Fig. 5E & Fig. S4**). However, in this case both responses remained elevated after the last stimulus in the train (probability of at least one spike in 50 ms after last pulse: mPFC = 0.38 ± 0.08, n = 12 cells / 4 mice; mTH = 0.44 ± 0.08, n = 14 cells / 7 mice) (**Fig. 5F**). Thus, despite their distinct short-term dynamics, NMDARs prominently contribute to the increase in evoked responses observed in both inputs. These findings indicate both cortical and thalamic inputs strongly impact CIN firing in the NAc core, with AMPARs mediating phasic responses and NMDARs having prolonged influences.

### Influence on ongoing action potential firing

Our initial experiments indicated that CINs in the NAc core exhibit low baseline firing rates at resting membrane potentials (**Fig. S4**), in line with previous reports of CINs in whole-cell recordings (Bennett and Wilson, 1999). However, CINs often display tonic firing *in vivo*, which could change the effects of cortical and thalamic inputs (Kimura et al., 1984; Aosaki et al., 1994; Bennett and Wilson, 1999; Sharott et al., 2012; Doig et al., 2014). To assess the influence of inputs on ongoing firing, we next repeated our current-clamp experiments in cells depolarized to maintain a baseline rate of 1-2 Hz (**Fig. 6A & S5**). Under these conditions, we found that both mPFC and mTH continued to activate CINs, with distinct dynamics (firing rate in 50 ms after first stimulus: mPFC = 15.3 ± 1.1, n = 23 cells / 14 mice; mTH = 7.0 ± 1.3, n = 18 cells / 12 mice; Mann Whitney test: U = 56; p < 0.0001) and persistence after stimulation (firing rate in 50 ms after last stimulus: mPFC = 19.7 ± 2.4; mTH = 19.7 ± 2.9; Mann Whitney test: U = 202.5; p = 0.91) (**Fig. 6B**). Lastly, additional recordings performed in the presence of selective antagonists indicated that both AMPARs and NMDARs continued to play influential roles (firing rate in 50 ms after first stimulus in CPP: mPFC = 10.8 ± 2.3, n = 11 cells / 8 mice; mTH = 5.4 ± 0.7, n = 11 cells / 6 mice; Mann Whitney test: U = 41; p = 0.21) (**Fig. 6C**), with the latter again contributing to the prolonged enhancement of firing (firing rate in 50 ms after last stimulus in NBQX: mPFC = 23.4 ± 5.0, n = 12 cells / 5 mice; mTH = 21.5 ± 4.9, n = 10 cells / 6 mice; Mann Whitney test: U = 56.5; p = 0.83) (**Fig. 6D**). Together, these findings demonstrate how mPFC and mTH can strongly influence action potential firing of CINs from either quiescent or spontaneously active states.

**Figure 6:**
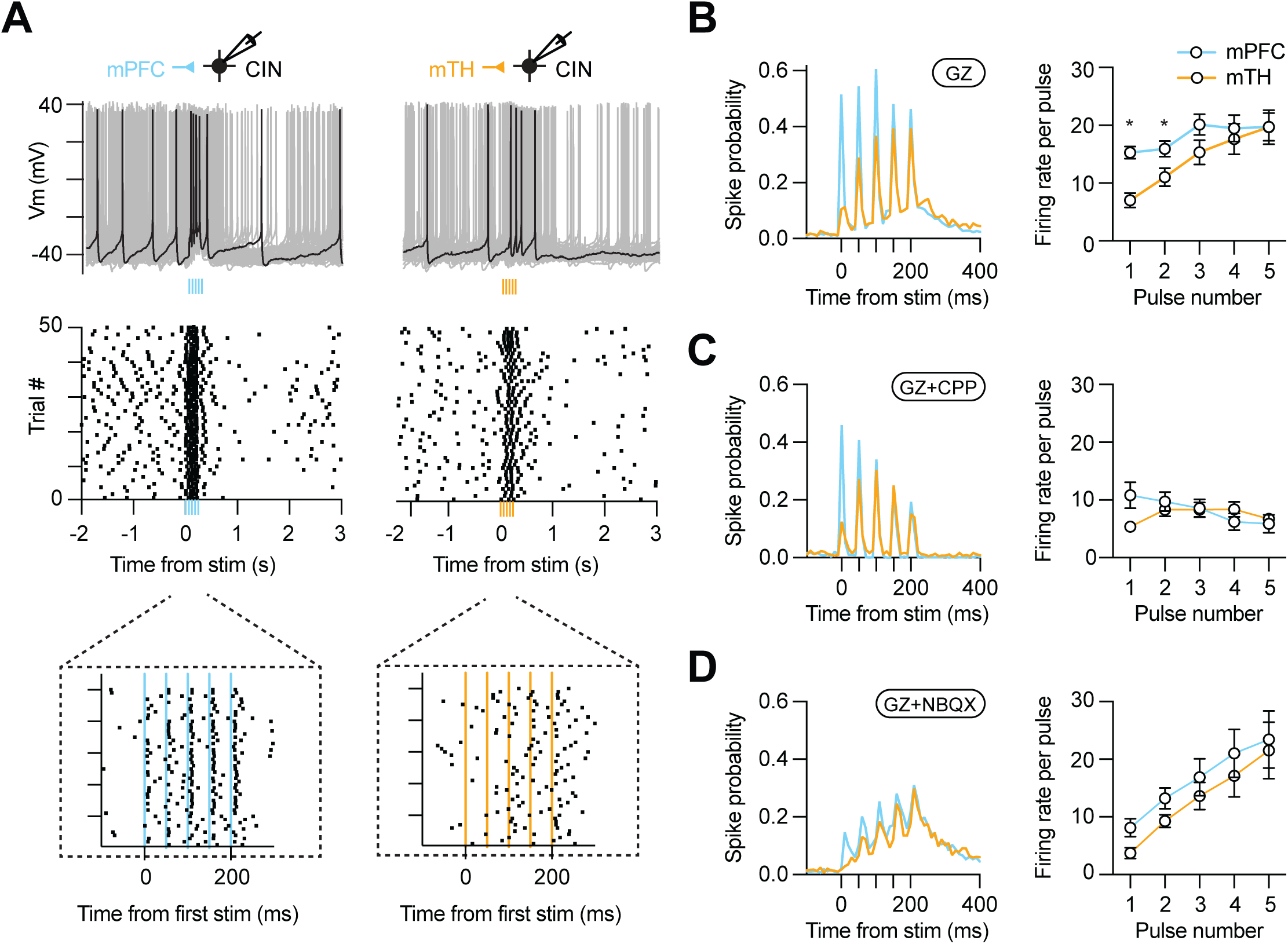
Influence on ongoing action potential firing. **(A)** *Top,* Examples of action potentials evoked by mPFC inputs (left) or mTH inputs (right) in current-clamp recordings from CINs depolarized to fire at 1-2 Hz. *Middle,* Raster plots of these recordings. *Bottom,* Expanded raster plots showing firing around stimulus pulses. **(B)** *Left,* Summary of spike probability versus time of stimulation for mPFC (blue, n = 23 cells / 14 mice) and mTH (orange, n = 18 cells / 12 mice) axons in the presence of GZ. Note that error bars are removed for clarity. *Right,* Summary of CIN activity calculated as firing rates following each stimulation pulse. **(C)** *Similar to (B)* with the addition of CPP to isolate AMPAR evoked spikes (mPFC, n = 11 cells / 8 mice; mTH, n = 11 cells / 6 mice). **(D)** Similar to *(B)* with the addition of NBQX to isolate NMDAR evoked spikes (mPFC, n = 12 cells / 5 mice; mTH, n = 10 cells / 6 mice).

## DISCUSSION

Here we examined how long-range inputs from prefrontal cortex and thalamus contact and influence CINs in the NAc core. We found these excitatory inputs exhibit distinct short-term dynamics, with depression at corticostriatal synapses and facilitation at thalamostriatal synapses. We also found these inputs engage AMPA and NMDA receptors at resting membrane potentials, with the latter having a prolonged influence on firing. Together, our findings demonstrate how CINs are driven by their excitatory inputs, with implications for control of signaling in the NAc core.

We first characterized the intrinsic properties of CINs in the NAc core, which we identified with Chat-eGFP mice as a small population of cells. We found CINs have spineless dendrites, which is similar to equivalent cells in the NAc medial shell (Baimel et al., 2022) and the dorsal striatum (Wilson et al., 1990). CINs *in vivo* often display tonic firing (Wilson et al., 1990; Doig et al., 2014), which can be paused by salient stimuli (Aosaki et al., 1994; Apicella et al., 2009), and is critical for behavior (Witten et al., 2010; Brown et al., 2012; Lee et al., 2016; Collins et al., 2019). Whether or not CINs also fire spontaneously *ex vivo* is variable, with some studies showing robust firing (Brown et al., 2012; Straub et al., 2014), and others relative quiescence (Bennett and Wilson, 1999). In our recording conditions, CINs in the NAc core displayed minimal spontaneous firing, similar to equivalent cells in the NAc medial shell (Baimel et al., 2022). Their evoked firing and intrinsic properties were similar to previous studies, including a h-current mediated voltage sag that can be important for shaping firing (Bennett et al., 2000; Straub et al., 2014).

Our initial anatomy identified inputs to CINs in the NAc core from across the brain using Cre-dependent rabies-mediated tracing (Callaway, 2008; Wall et al., 2010). Previous studies have used this approach to examine inputs to projection neurons and interneurons in both the dorsal striatum (Wall et al., 2013; Guo et al., 2015; Klug et al., 2018) and NAc (Li et al., 2018; Baimel et al., 2022). In the dorsal striatum, cortical inputs to CINs primarily arise from motor and somatosensory areas (Doig et al., 2014; Mamaligas et al., 2019; Johansson and Silberberg, 2020), whereas CINs in the NAc medial shell receive relatively few cortical inputs (Baimel et al., 2022). In contrast, we found that excitatory equivalent inputs to the NAc core primarily originate from the prefrontal cortex. In the dorsal striatum, thalamic inputs to CINs largely arise from intralaminar nuclei, including the parafascicular nucleus (Lapper and Bolam, 1992; Guo et al., 2015; Johansson and Silberberg, 2020), whereas CINs in the NAc medial shell receive inputs from the paraventricular nucleus (Baimel et al., 2022). In contrast, we found the NAc core receives inputs from several nuclei in medial and midline thalamus, with few inputs from the intralaminar nuclei. These results highlight how CINs in different regions of the striatum, and even within different subregions of the NAc, receive unique distributions of cortical and thalamic inputs.

Although the basolateral amygdala is a major input to the NAc core (Stuber et al., 2011; Britt et al., 2012; MacAskill et al., 2012), it was sparsely labeled in our rabies experiments, similar to our recent findings in the NAc medial shell (Baimel et al., 2022). This likely does not reflect virus tropism, because rabies tracing in D1-Cre and D2-Cre mice of the NAc core labels amygdala inputs (Li et al., 2018). Another notable absence in our data is inputs from the ventral tegmental area, despite known synapses onto CINs in the NAc core (Brown et al., 2012; Chuhma et al., 2014). It is possible these synapses are poorly labeled by Cre-dependent rabies tracing methods due to how the virus is taken up by dopaminergic synapses (Wall et al., 2013). Finally, we also observed prominent inputs from pallidal areas, particularly the ventral pallidum, which are thought to make predominantly inhibitory GABA-mediated projections to the NAc (Vachez et al., 2021).

Our subsequent slice physiology examined the sign, dynamics, and receptor contributions of cortical and thalamic inputs. We found that both inputs are excitatory at CINs, with no mono-synaptically evoked GABA-A receptor mediated IPSCs. Interestingly, both cortical and thalamic inputs were larger onto MSNs, consistent with engagement of multiple cell types in the circuit. In other parts of the NAc, long-range excitatory inputs can engage GABAergic interneurons, which along with MSNs can mediate feed-forward inhibition (Scudder et al., 2018). While we did not test it here, in the NAc medial shell, these polysynaptic networks also influence the firing properties of CINs (Baimel et al., 2022). Moreover, the activation of CINs can feedback onto GABAergic interneurons and MSNs to shape network activity (English et al., 2012; Dorst et al., 2020; Kocaturk et al., 2022). In the future, it will be interesting to examine how cortical and thalamic inputs engage additional cell types in the NAc core to generate these more complex polysynaptic responses.

During repetitive activity, we found that cortical and thalamic inputs exhibit distinct synaptic dynamics. Previous studies in the dorsal striatum indicate that parafascicular thalamic inputs display robust facilitation (Ding et al., 2010; Doig et al., 2014; Johansson and Silberberg, 2020). Similar to these results, in the NAc core, we find that medial and midline thalamic inputs also evoke robust facilitation at CINs. In contrast, previous work in dorsal striatum has been mixed for cortical inputs, which are sometimes weak and can depress or facilitate (Ding et al., 2010; Doig et al., 2014; Johansson and Silberberg, 2020). In the NAc core, we find that cortical inputs evoke strong responses that are maintained over trains but display some depression at CINs. Interestingly, during more sustained activity, the currents evoked by cortical and thalamic inputs approach similar levels, which suggested they can both influence action potential firing.

Selective pharmacology allowed us to dissociate the roles of AMPA and NMDA receptors to synaptic responses. We found that AMPA receptors make strong contributions at both inputs, highlighting differences in synaptic dynamics. Previous results indicate that NMDA receptors are primarily activated by thalamic inputs, with relatively little engagement by cortical inputs (Kosillo et al., 2016; Mamaligas et al., 2019). In contrast, we found that NMDA receptors also make strong contributions at both inputs, which are readily engaged even at resting potentials of -50 mV. As expected due to their slow kinetics (Lester et al., 1990), NMDA receptors responses integrate, with synaptic charge building up during repetitive activity. Interestingly, despite their distinct synaptic dynamics, this NMDA receptor charge is similar at cortical and thalamic inputs.

To assess impact on action potential firing, we first examined CINs held at a quiescent state and found that cortical and thalamic inputs have distinct effects. Consistent with their different dynamics, cortical inputs evoked an initial strong response, which diminished over trains due to depression. In contrast, thalamic inputs evoked an initial weak response on firing, which built over trains due to facilitation. Also consistent with our voltage-clamp recordings, both AMPA and NMDA receptors play a role in this evoked firing. AMPA receptor responses were transient for both inputs, and unable to drive sustained firing on their own. NMDA receptor responses are sustained, continuing to build during repetitive activity, and ultimately making a major contribution to firing, with action potentials less time locked to stimuli. Thus, in contrast to related circuits in the dorsal striatum (Kosillo et al., 2016; Mamaligas et al., 2019), NMDA receptors play a key role in CINs in the NAc core, allowing sustained responses to both cortical and thalamic inputs.

We observed similar results in CINs that were depolarized in order to maintain spontaneous action potential firing. As in quiescent cells, cortical inputs were initially stronger, thalamic inputs facilitated to drive activity, and NMDA receptor contributions grew during trains. In both cases, we also observed reduced spontaneous firing after the stimulus, which could be due to a variety of factors. For example, in the dorsal striatum, high frequency firing can engage intrinsic conductances that shut down CIN firing for prolonged periods (Zhang et al., 2018). In contrast, in the NAc medial shell, activation of GABAergic interneurons can robustly pause CIN firing (Baimel et al., 2022), although this is unlikely here, as inhibition was blocked. Lastly, activation of CINs triggers the release of acetylcholine, which can in turn evoke presynaptic dopamine or GABA release (Cachope et al., 2012; Threlfell et al., 2012; Nelson et al., 2014; Liu et al., 2022), that can feedback to inhibit CIN firing (Chuhma et al., 2014; Straub et al., 2014; Wieland et al., 2014). In the future, it will be interesting to examine which mechanisms cause the pause in the NAc core.

In summary, we have characterized how CINs in the NAc core receive and process cortical and thalamic inputs. Inputs from medial and midline thalamus are prominent, with robust facilitation and activation of both AMPA and NMDA receptors to influence firing. Inputs from prefrontal cortex are also efficacious, with distinct synaptic dynamics but broadly similar effects on action potential firing. Our results contrast with the long-standing model from dorsal striatum that CINs are primarily driven by thalamic inputs. In the future, it will be interesting to consider additional long-range inputs, including from ventral pallidum, which may inhibit CINs. Lastly, our findings highlight how different subregions and cell types in the NAc receive and process distinct synaptic inputs, which are ultimately responsible for generating network activity that shapes animal behavior.

## Acknowledgements

We thank the Carter lab for helpful discussions and comments on the manuscript. This work was supported by NIH F30 MH129055 (EVJ), NIH R01 MH085974 (AGC), and NIH R21 DA064023 (AGC). The authors have no financial conflicts of interest.

---

**Figure S1:**
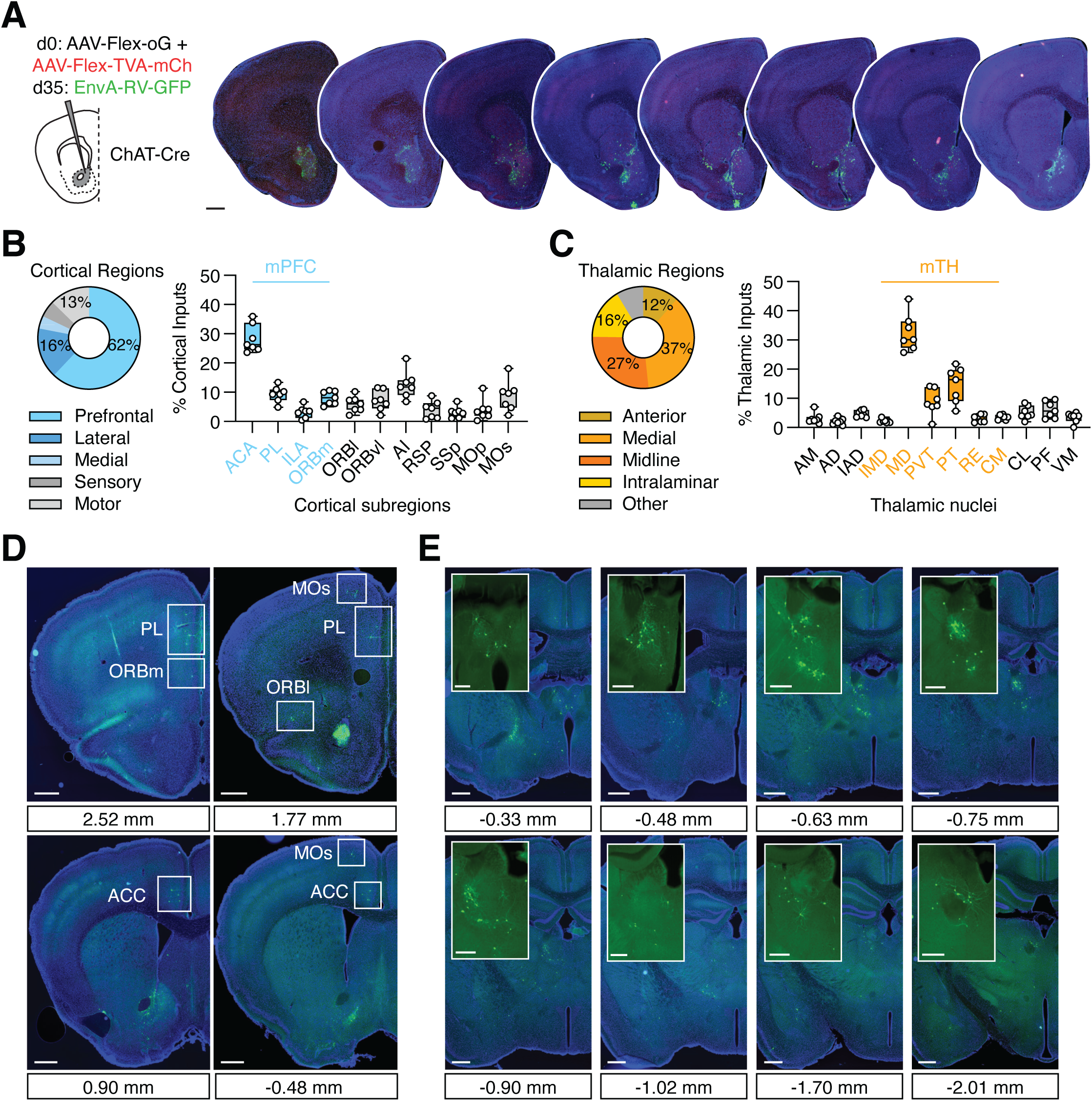
Additional analysis of rabies anatomy. **(A)** *Left,* Schematic of injection strategy, similar to Figure 2. *Right,* Images of coronal slices from ChAT-Cre animal, including the injection site and local monosynaptic inputs to CINs in the NAc core. Scale bar = 500 µm. **(B)** *Left,* Breakdown of cortical inputs by region across animals (n = 7). *Right,* Breakdown of cortical subregions with medial prefrontal cortex (mPFC) shown on x-axis in blue. **(C)** *Left,* Breakdown of thalamic inputs by subregion (n = 7). *Right,* Breakdown of nuclei in medial and midline regions (mTH) shown on x-axis in orange. **(D)** Example images of select cortical areas, including prelimbic (PL), orbitofrontal (ORB), anterior cingulate (ACC), and secondary motor (MOs) cortex. Scale bar = 500 µm. **(E)** Example images of thalamus across anterior-posterior axis. Scale bar = 500 µm. Inset scale bar = 250 µm.

**Figure S2:**
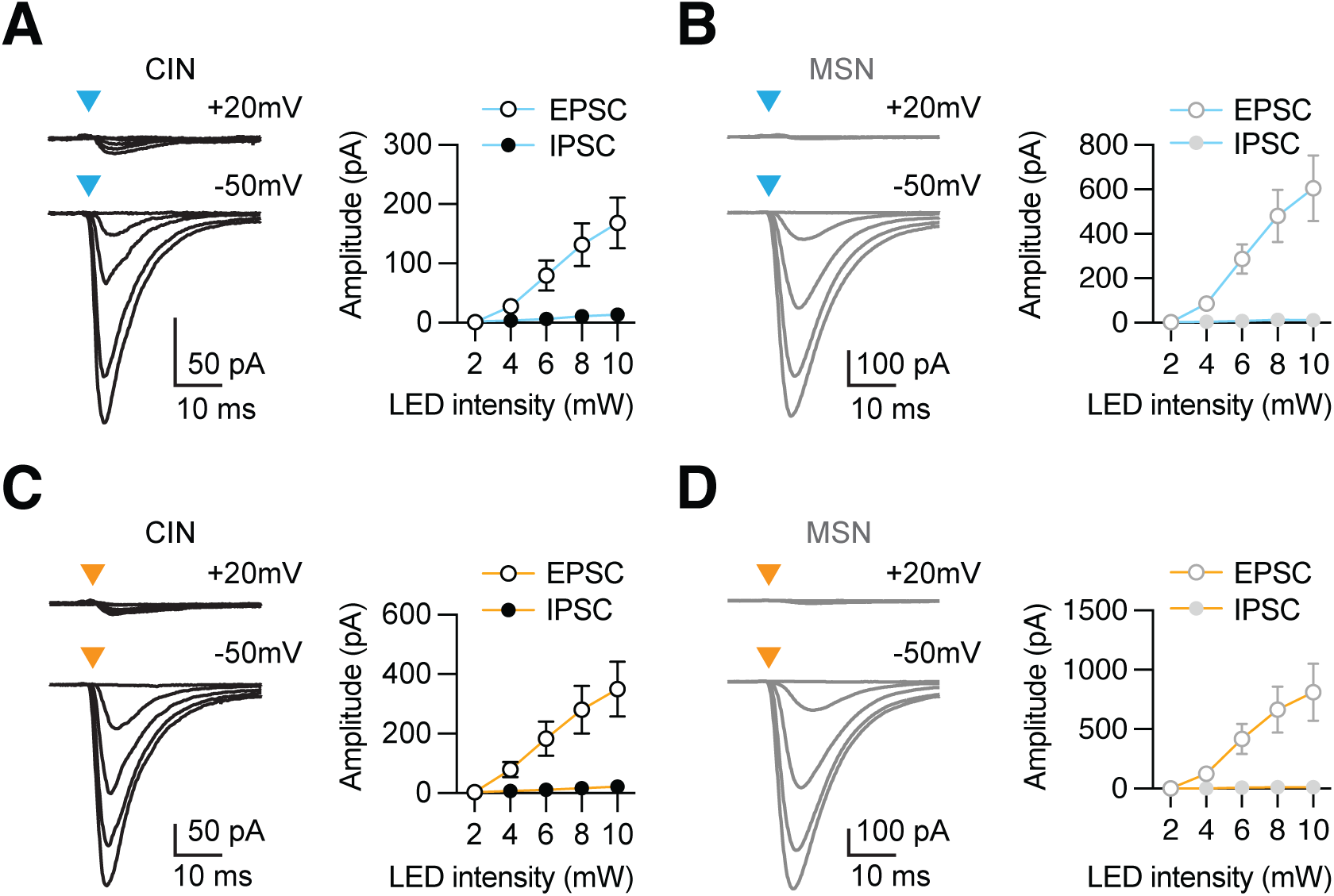
Further comparison of synaptic responses at CINs and MSNs. **(A)***Left,* Average mPFC-evoked EPSCs and IPSCs recorded at -50 mV (E_GABA_) and +20 mV (E_AMPA_), respectively, across five stimulus intensities in CINs (n = 9 cells / 5 mice). *Right,* Summary of EPSC and IPSC amplitudes across stimulus intensities. **(B)** Similar for mPFC inputs to MSNs in cell pairs. **(C)** Similar to (A) for mTH inputs to CINs (n = 8 pairs / 3 mice). **(D)** Similar to (B) for mTH inputs to MSNs in cell pairs.

**Figure S3:**
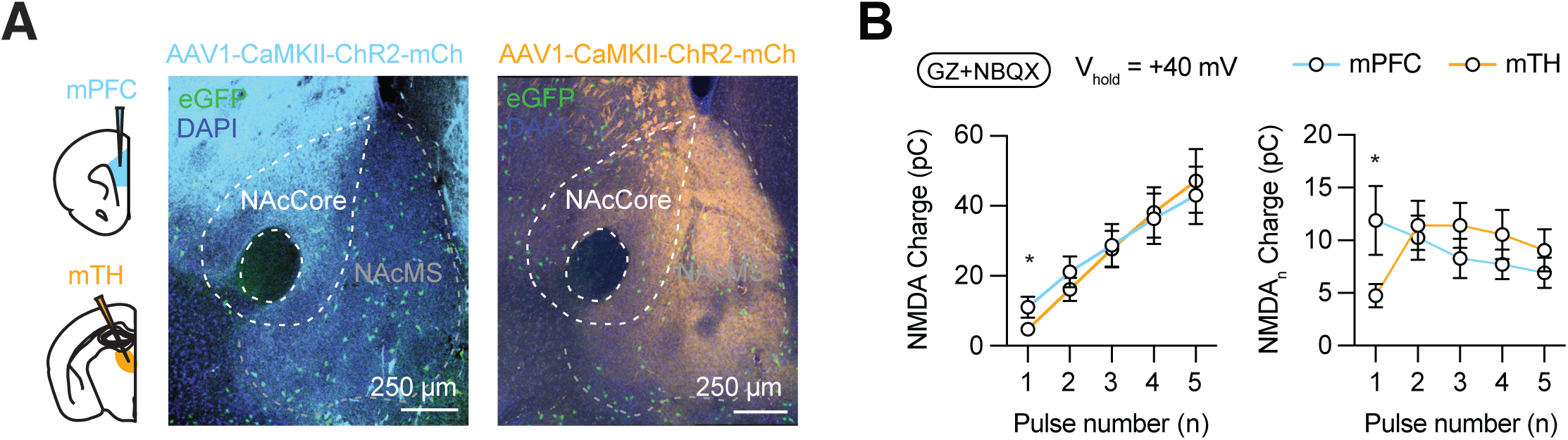
Supplemental analysis of responses to stimulus trains. **(A)** *Left,* Schematic of AAV1-CaMKII-ChR2-mCherry injections into mPFC or mTH of ChAT-eGFP mice. *Right*, Image of mPFC axons (blue) or mTH axons (orange) in the NAc core and neighboring NAc medial shell (NAcMS) for comparison. **(B)** *Left,* Summary of cumulative charge of mPFC (blue) and mTH (orange) evoked NMDAR-mediated EPSCs in CINs recorded at +40 mV. *Right,* Charge as a function of pulse number (n), showing the charge evoked by the addition of each subsequent pulse.

**Figure S4:**
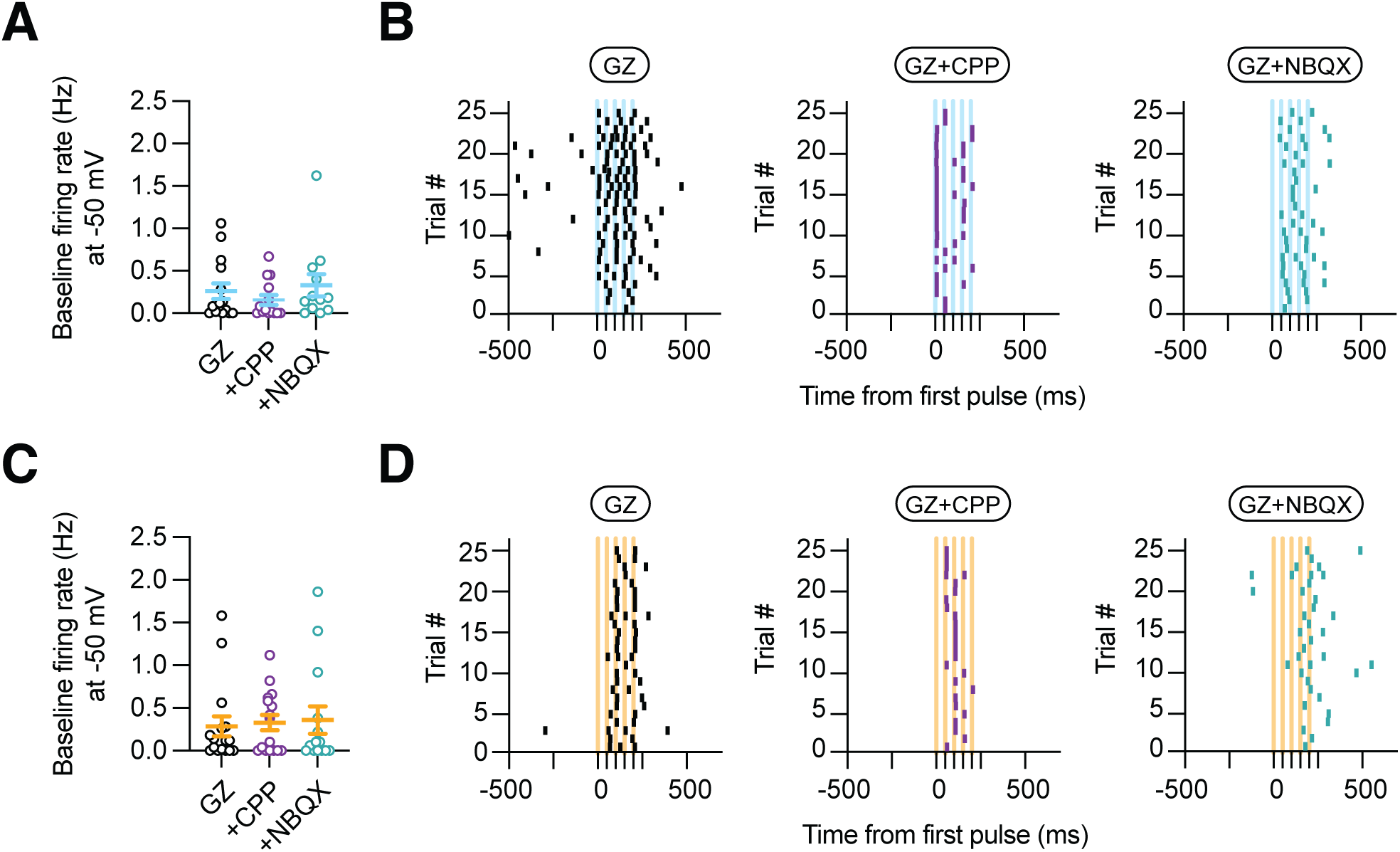
Additional analysis of evoked spiking from quiescent CINs. **(A)** Summary baseline firing rates of CINs held near -50 mV and responding to mPFC inputs in the presence of GZ (0.26 ± 0.09 Hz, n = 15 cells / 8 mice), GZ and CPP (0.16 ± 0.06 Hz, n = 14 cells / 6 mice), or GZ and NBQX (0.34 ± 0.15 Hz, n = 12 cells / 4 mice). **(B)** Example raster plots of CINs responding to mPFC inputs in the presence of GZ (left), GZ and CPP (middle), or GZ and NBQX (right), showing 500 ms before and 500 ms after stimulation (blue lines). **(C-D)** Similar to (A-B) for CINs responding to mTH inputs in the presence of GZ (0.29 ± 0.12 Hz, n = 16 cells / 8 mice), GZ and CPP (0.33 ± 0.09 Hz, n = 16 cells / 5 mice), or GZ and NBQX (0.36 ± 0.16 Hz, n = 14 cells / 7 mice).

**Figure S5:**
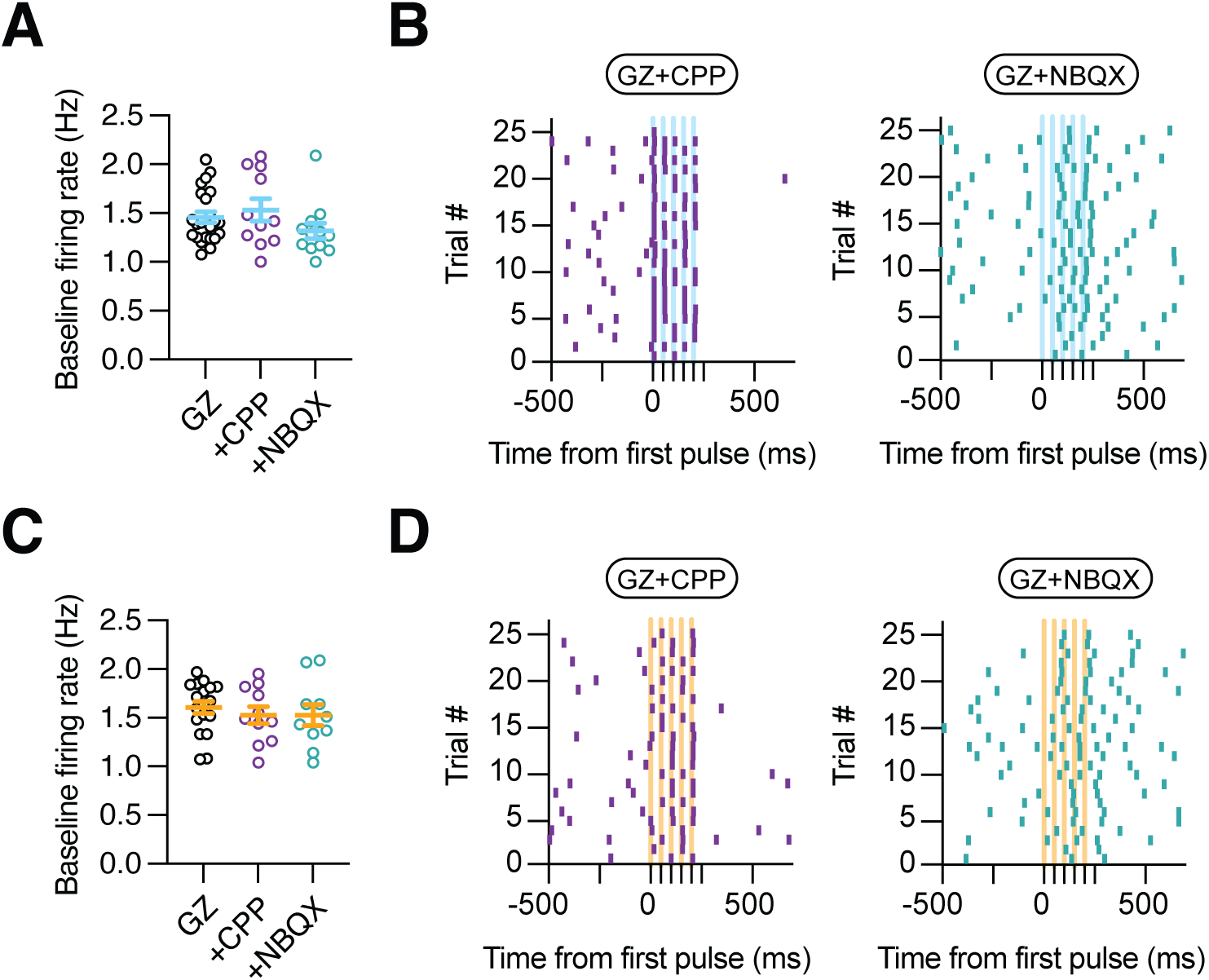
Further analysis of evoked spiking in spontaneously active CINs. **(A)** Summary baseline firing rates of CINs firing between 1-2 Hz and responding to mPFC inputs in the presence of GZ (1.46 ± 0.06 Hz, n = 23 cells / 14 mice), GZ and CPP (1.53 ± 0.11 Hz, n = 11 cells / 8 mice), or GZ and NBQX (1.32 ± 0.08 Hz, n = 12 cells / 5 mice). **(B)** Example raster plots of CINs responding to mPFC inputs in the presence of GZ and CPP (left) or GZ and NBQX (right), showing 500 ms before and 500 ms after stimulation (blue lines). **(C-D)** Similar to (A-B) for CINs responding to mTH inputs in the presence of GZ (1.61 ± 0.06 Hz, n = 18 cells / 12 mice), GZ and CPP (1.53 ± 0.09 Hz, n = 11 cells / 6 mice), or GZ and NBQX (1.53 ± 0.11 Hz, n = 10 cells / 6 mice).

## Notes

### Competing Interest Statement

The authors have declared no competing interest.

